# A previously established divergent lineage of the hybrid fungal pathogen *Verticillium longisporum* emerges as stem striping pathogen in British oilseed rape

**DOI:** 10.1101/102541

**Authors:** Jasper R. L. Depotter, Michael F. Seidl, Grardy C. M. van den Berg, Bart P.H.J. Thomma, Thomas A. Wood

**Author notes:** These authors contributed equally (Jasper R. L. Depotter, Michael F. Seidl). These authors contributed equally (Bart P.H.J. Thomma, Thomas A. Wood). For correspondence: Bart P.H.J. Thomma, Laboratory of Phytopathology, Wageningen University, Droevendaalsesteeg 1, 6708 PB Wageningen, The Netherlands.

## Abstract

Population genetic structures illustrate evolutionary trajectories of organisms adapting to differential environmental conditions. Pathogen populations are typically shaped by co-evolution with their hosts through genetic co-structuring. Verticillium stem striping was mainly observed in continental Europe, but has recently emerged in other countries including the United Kingdom. The disease is caused by the hybrid fungal species *Verticillium longisporum* that originates from at least three separate hybridization events, yet strains from the hybridization event between *Verticillium* progenitor species A1 and D1 are mainly responsible for Verticillium stem striping. By using multi-locus genotype analysis, we reveal a hitherto un-described dichotomy within *V. longisporum* lineage A1/D1 that correlates with the geographic distribution of the isolates with an “A1/D1 West” and an “A1/D1 East” cluster according to their relative location in Europe. Genome comparison between representatives of the A1/D1 West and East clusters confirmed mutual common origin, excluding distinctiveness through separate hybridization events. The A1/D1 West population is responsible for the sudden emergence of Verticillium stem striping in the UK. Remarkably, this emergence is caused by a British *V. longisporum* population that is genetically more diverse than the entire A1/D1 East cluster. Conceivably, *V. longisporum* has previously established in the UK, but remained latent or undiagnosed as an oilseed rape pathogen until recently. This finding illustrates that a recent introduction is not a prerequisite for a pathogen to emerge, as environmental factors and cultural practices can also play a pivotal role in outbreaks of novel diseases.

## Introduction

Interspecific hybridization, the natural or induced combination of two genetically divergent parents, is pervasive among many different eukarotic taxa such as plants, insects, birds, mammals and fungi (Brasier 2000; Mallet 2005). Hybrids that receive a genome copy of both parents initially double their chromosome number, and experience a so-called “genome shock” that incites major genomic reorganizations that can manifest by genome rearrangements, extensive gene loss, transposon activation, or alterations in gene expression (Doyle *et al.* 2008). Due to this increased genome plasticity, hybridization can promote the emergence of altered phenotypes that allow adaptation to novel niches or to changing environments. Consequently, many fungal hybrids are exploited in the production of food and beverages. For example, at least two recent hybridization events between the well-known brewing yeast *Saccharomyces cerevisiae* and its close relative *Saccharomyces eubayanus* gave rise to *Saccharomyces pastorianus*, a lineage with high cold tolerance and good maltose/maltotriose utilization capabilities (Gibson & Liti 2015). Both characteristics are exploited in the production of the popular lager beer that is generated from malted barley at very low temperatures. Similarly, interspecific hybridization is also a potent driver for the evolution of fungal plant pathogens as increased genome plasticity promoted by an hybridization event allows hybrids to differentiate and acquire new pathogenic traits (Depotter *et al.* 2016b).

*Verticillium* species are causal agents of wilt diseases on many economically important crops, with a total estimated annual loss of €3 billion worldwide in the 20 most affected hosts, including cotton and olive (Depotter *et al.* 2016a). *Verticillium dahliae* is the most notorious wilt agent of this genus and is characterized by its extremely broad host range that encompasses hundreds of hosts (Fradin & Thomma 2006). *V. dahliae* propagates asexually and genomic variation is established by mechanisms different from meiotic recombination, such as large-scale genomic rearrangements, horizontal gene transfer and transposon activity (de Jonge *et al.* 2012, 2013; Seidl & Thomma 2014; Faino *et al.* 2016). Moreover, *Verticillium* experienced more intrusive genomic evolution by inter-specific crosses within the genus leading to an approximate doubling of the genome size. Interspecific *Verticillium* hybrids gave rise to new diseases such as Verticillium stem striping on oilseed rape (Inderbitzin *et al.* 2011b; Depotter *et al.* 2016a). At least three hybridization events between two separate *Verticillium* spp. have occurred that have been classified under the same species name, *V. longisporum* (Karapapa *et al.* 1997; Inderbitzin *et al.* 2011b). The three hybrid lineages have been named after their respective hybridization parents: A1/D1, A1/D2 and A1/D3 (Inderbitzin *et al.* 2011b). A1 and D1 are hitherto un-described *Verticillium* species, whereas D2 and D3 are presumed *V. dahliae* isolates. Similar to other hybrid pathogens, hybridization appears to have altered the host range of *Verticillium* (Depotter *et al.* 2016b). *V. longisporum* is highly adapted to brassicaceous hosts, such as oilseed rape and cauliflower, whereas *V. dahliae* generally does not colonize these plants (Depotter *et al.* 2016a). Moreover, differences in pathogenicity are also observed between hybrid lineages, as *V. longisporum* A1/D1 and A1/D3 are often found on multiple brassicaceous hosts, whereas lineage A1/D2 has hitherto only been found on horseradish (Inderbitzin *et al.* 2011b; Yu *et al.* 2016). Furthermore, lineage A1/D1 is predominantly found on oilseed rape and responsible for the Verticillium stem striping disease as this lineage is the most virulent on this crop (Novakazi *et al.* 2015).

Verticillium stem striping is a relatively new disease that was first reported in the west and south of Scania, southern Sweden, in 1969 (Kroeker 1970). Until recently, this oilseed rape disease was only present in North-central Europe, but over the last decade other important oilseed rape production regions have also been affected (Gladders *et al.* 2011; CFIA 2015). Verticillium stem striping was noticed in UK oilseed rape production for the first time in 2007, in the counties Kent and Herefordshire (Gladders *et al.* 2011). Currently, *V. longisporum* is present throughout England, yet the disease is most prevalent in the east (Gladders *et al.* 2013). The main causal agent of Verticillium stem striping, lineage A1/D1, has also been found outside Europe in Japan and the USA, albeit on different crops than oilseed rape (Carder & Barbara 1994; Subbarao *et al.* 1995). The wide-spread occurance of *V. longisporum* A1/D1 indicates the importance of human activity in the spread of this disease as *V. longisporum* is concidered soil-borne without the long-distance dispersal of air-borne spores (Depotter *et al.* 2016a). Dispersal of Verticillium by the trade of plant commodities has been observed for *V. dahliae*, which has facilitated the intercontinental spread of the pathogen (Atallah *et al.* 2012).

The aim of this study is to reveal population structures within Verticillium stem striping lineage A1/D1 by screening of rapidly evolving DNA regions. Microsatellites or simple sequence repeat (SSR) loci are hyper-variable genome regions of simple DNA motifs repeated in tandem. The repetitive character of these regions makes them more prone to mutation than non-repetitive sequences due to unequal crossing-over and replication slippage, generally revealing high degrees of polymorphism (Levinson & Gutman 1987). Several population studies previously used SSR loci to described diversity within *Verticillium* populations (Atallah *et al.* 2010, 2012). However, hitherto, no population studies have been performed on *V. longisporum*. Here, we assessed the genetic diversity within a broad geographic range of the *V. longisporum* lineage A1/D1 isolates and, surprisingly, found clear population structuring. The origin of these distinct populations was further elucidated by genealogical analysis and genome comparison. Here, established Verticillium stem striping populations (e.g. from Germany and Sweden) were compared with populations from a recent disease outbreak in the UK, in order to link the population dynamics with the expansion of Verticillium stem striping.

## Materials and Methods

### Isolate collection, DNA extraction and lineage characterization

In total 88 isolates were collected from nine different countries (Table S1). The UK and Latvian isolates were isolated form diseased oilseed rape stems that were attained by the National Institute of Agricultural Botany (NIAB, Cambridge, UK) and Integrētās Audzēšanas Skola (SIA, Riga, Latvia), respectively. All other isolates used in this study were obtained from the donors as mentioned in Table S1. The lineage and mating type to which the respective isolates belong was determined previously for several isolates (Table S1; Inderbitzin *et al.*, 2011; Inderbitzin *et al.*, 2013). Mycelium was harvested from two-week-old potato dextrose broth cultures and DNA was extracted according to DNA extraction protocol A from Ribeiro and Lovato (2007). DNA suspensions were stored at -20°C until use.

The hitherto uncharacterized isolates were screened for the presence of a marker that is specific for lineage A1/D1. To this end, the primer pair D1f/AlfD1r was used for PCR to amplify a fragment from the *GPD* locus according to Inderbitzin *et al.* (2013). Amplicons were displayed by gel electrophoresis on a 1% agarose gel. Furthermore, isolates were screened for *MAT1-1* and *MAT1-2* idiomorphs with the primer pairs Alf/MAT11r and HMG21f/MAT21r, respectively, according to Inderbitzin *et al.*, (2011b). Amplicons were displayed by gel electrophoresis on a 1.5% agarose gel.

### SSR loci

A genome wide screening for polymorphic SSR loci was done with unpublished draft genome sequences from several *V. longisporum* isolates. INDELs between genomes were extracted from the whole-genome alignments using the mummer package (v3.1) (Kurtz *et al.* 2004). Gene sequence variations were received by the variant call format tool (Danecek *et al.* 2011). Insertions and deletions between 5 and 20 nucleotides were selected and screened for recurrent patterns that are typical for SSR loci. Primers were developed for 61 putative polymorphic SSR loci with the Primer3 software (Untergasser *et al.* 2012).

Additional polymorphic SSR markers for lineage A1/D1 were used in this study. VD1, VD8 and VD12 from Atallah *et al.* (2010) were originally designed for *V. dahliae* and were found to be polymorphic between *V. longisporum* isolates. In addition, VDA783, VDA787 and VDA823 from Barbara *et al.* (2005), designed for *V. longisporum*, were used.

SSR loci were labelled and amplified with an M13 fluorescent tag according to Schuelke (2000). The PCRs consisted of a 2 min initial denaturation step at 95°C, 30 cycles of 35 sec at 95°C, 45 sec at 62°C, and 1 min at 72°C, followed by 8 cycles of 30 sec at 95°C, 45 sec at 53°C and 1 min at 72°C, followed by an extension of 10 min at 72°C. The PCR mix contained 8 pmol of each reverse and M13 tagged universal sequence primer and 2.5 pmol of the forward primer in a final 10μl reaction volume: 1X GoTaq^®^ Flexi Buffer Mg-free, 2.5mM MgCl_2_, 0.2mM each dNTP, 100ng template DNA, DNA polymerase, 1.25 u GoTaq^®^ polymerase (Promega, Madison, WI, USA). The labelled PCR products were then combined with Hi-Di formamide and LIZ-500 size standard and resolved on a 3730XL DNA Analyzer (Applied Biosystems, Foster City, CA, USA). The results were processed using GeneMapper v4.0 software (Applied Biosystems, Foster City, CA, USA).

### Population structure

*V. longisporum* isolates were individually clustered based on polymorphic SSR loci using the software Structure version 2.3 (Pritchard *et al.* 2000). Despite the amphidiploid character of *V. longisporum*, the data was analyzed as for a haploid organism as all markers only gave a single polymorphic signal. The population was tested for containing 1 up to 6 genetic clusters (K). For every cluster, 10 runs were performed with a burn-in period of 500,000 generations and 1,000,000 Markov Chain Monte Carlo (MCMC) simulations. The admixture model was chosen and the loci were considered independent. The most likely number of genetic clusters in the population was determined with the ad-hoc statistic Δ*K* (Evanno *et al.* 2005) using Structure Harvester (Earl & vonHoldt 2012). Hereby, the amount of clusters in a population is determined based on the rate of change in the log probability of data between successive K values. Furthermore, the results from Structure were permuted and aligned in the program CLUMPP 1.1.2 (method Greedy, random input order, 1,000,000 repeats) (Jakobsson & Rosenberg 2007) and visualized with the software Distruct 1.1 (Rosenberg 2004). The correlation between the clusters and the country of origin of the isolates was determined with the Spearman’s rank correlation coefficient (*ρ*) with Hmisc package in R 3.2.3 (R Core Team 2015; Harrell & Dupont 2016).

The standardized index of association *I*_A_^S^ gives an indication for the recombination rate between organisms. In outcrossing populations, no linkage disequilibrium is present and an *I*_A_^S^ of 0 is expected (Burt *et al.* 1996). *I*_A_^S^ was computed to test the recombination potential within the *V. longisporum* lineage A1/D1 population using the software LIAN version 3.7 (Haubold & Hudson 2000). A Monte Carlo simulation of 100,000 iterations was chosen.

The genetic clusters identified by Structure were evaluated using an analysis of molecular variance (AMOVA) with the software GenAlEx version 6.502 (Peakall & Smouse 2006, 2012). The variance within the genetic clusters was compared was with the variance between the genetic clusters with an analogue of Wright’s fixation index (*Φ*_*PT*_) (Excoffier *et al.* 1992).

The population diversity was assessed for populations with a minimum of 10 representatives, excluding isolates with missing data. Multi-locus genotype (MLG) diversity and the allelic richness (*AR*) were determined for each population using software Contrib 1.4, using rarefaction size 5 (Petit *et al.* 1998). Nei’s (1973) genetic diversity corrected for sample size (*H*_*s*_) values were generated in GenoDive (Meirmans & Van Tienderen 2004).

Genealogical relationships among the different MLGs haplotypes in the *V. longisporum* population was inferred using the median-joining method (Bandelt *et al.* 1999), implemented in the software Network 5.0.0.0 (http://www.fluxus-engineering.com). All SSR loci were weighed equally (10) and an epsilon = 0 was chosen. The hypervariable SSR locus VD12 was not included in the analysis to reduce the amount of MLGs, and MLGs with missing data were excluded as well.

### Genome sequencing and assembly of two *Verticillium longisporum* isolates

Genomic DNA of *V. longisporum* isolates VL20 and VLB2 was isolated from conidia and mycelium fragments that were harvested from 10-day-old cultures grown in liquid potato dextrose agar according to the protocol described by Seidl *et al.* (2015). The PacBio libraries for sequencing on the PacBio RSII machine (Pacific Biosciences of California, CA, USA) were constructed using ~20 μg of *V. longisporum* DNA, similary as described previously by Faino *et al.*, (2015). Briefly, DNA was mechanically sheared, and size selected using the BluePippin preparation system (Sage Science, Beverly, MA, USA) to produce ~20 kb insert size libraries. The sheared DNA and final library were characterized for size distribution using an Agilent Bioanalyzer 2100 (Agilent Technology, Inc., Santa Clara, CA, USA). The PacBio libraries were sequenced on six SMRT cells per *V. longisporum* isolate at KeyGene N. V. (Wageningen, the Netherlands) using the PacBio RS II instrument. Sequencing was performed using the P6-C4 polymerase-Chemistry combination and a >4h movie time and stage start. Post filtering, a total of 344,906 (N50 ~20 kb) and 358,083 (N50 ~19 kb) polymerase reads were obtained for *V. longisporum* isolates VLB2 and VL20, respectively. Filtered sub-reads for VLB2 and VL20 (457,986; N50 ~14 kb and 466,673; N50 ~13.7 kb, respectively) were assembled using the HGAP v3 protocol (Chin *et al.* 2013). Subsequently, the HGAP3 assemblies underwent additional polishing using Quiver (Chin *et al.* 2013). The *de novo* assemblies were further upgraded using FinisherSC, and the upgraded assemblies were polished with Quiver (Lam *et al.* 2015). Lastly, contigs that displayed very low or exceptionally high PacBio read coverage, as well as the contig representing the mitochondrial DNA, were removed from the final assemblies. *De novo* repetitive elements in the genomes of *V. longisporum* were identified using RepeatModeler (v1.0.8), and repetitive elements were subsequently masked using RepeatMasker (v4.0.6; sensitive mode).

### Genome comparisons between *V. longisporum* and *V. dahliae* strains

To place the newly sequenced *V. longisporum* isolates in context of 74 previously analysed *Verticillium* spp., we identified the alleles (A or D) of four previously used protein-coding genes actin (*ACT*), elongation factor 1-alpha (*EF*), glyceraldehyde-3-phosphate dehydrogenase (*GPD*) and tryptophan synthase (*TS*) in the genome assemblies of VLB20 and VLB2 using blastn searches (Inderbitzin *et al.* 2011a). Sequences were extracted from the genome assemblies and aligned to the four genes in the 74 *Verticillium* isolates using mafft (LINSi; v7.271) (Katoh & Standley 2013). A phylogenetic tree was reconstructed using PhyML using the GTR nucleotide substitution model and four discrete gamma categories (Guindon & Gascuel 2003). The robustness of the phylogeny was assessed using 500 bootstrap replicates.

Whole-genome comparisons between *V. longisporum* strains VLB2 and VL20 and between *V. dahliae* strains JR2 and VdLs17 (Faino *et al.* 2015) were performed with nucmer (maxmatch), which is part of the mummer package (v3.1) (Kurtz *et al.* 2004). Small polymorphisms (SNPs and INDELs) between genomes were extracted from the whole-genome alignments using show-snps (excluding ambiguous alignments), which is part of the mummer package (v3.1). Lineage-specific regions per individual *Verticillium* strain were determined by extracting nucmer alignments and subsequently identifying genomic regions that lack alignments with the other isolate (bedtools genomecov) (Quinlan & Hall 2010).

To tentatively assign the individual sub-genomes, the repeat-masked genome of *V.dahliae* strain JR2 was compared to the repeat-masked *V. longisporum* genomes using nucmer (maxmatch), of which only 1-to-1 alignments longer than 5 kb where retained. Genomic regions were assigned to parental sub-genomes based on the average identity of consecutive alignments (defined by location and/or strand), where regions with average identity >95% were assigned to D and < 95% identity to A, respectively. The pairwise identity between A and D parents within and between *V. longisporum* strains was calculated using nucmer (mum), with dividing the respective query sequences into non-overlapping windows of 500 bp.

## Results

A geographically diverse collection of *V. longisporum* isolates was obtained in order to assess the population diversity and to genotypically link isolates from different origins. Isolates from 9 different countries were included in this study: UK (*n* = 25), Germany (*n* = 22), Sweden (*n =* 15), USA (*n* = 6), Japan (*n* = 6), France (*n* = 6), Belgium (*n* = 5), the Netherlands (*n* = 2) and Latvia (*n* = 1) (Table S1). In total, 88 *V. longisporum* isolates were screened for SSR polymorphisms in order to determine the population structure. In total, 61 putative polymorphic SSR markers were developed based on unpublished draft *V. longisporum* genome sequence data and tested. Nine markers displayed polymorphisms in the tested *V. longisporum* collection (Table 1). Our analysis was performed as for a haploid organism as all markers only gave a single polymorphic signal.

**Table 1:**
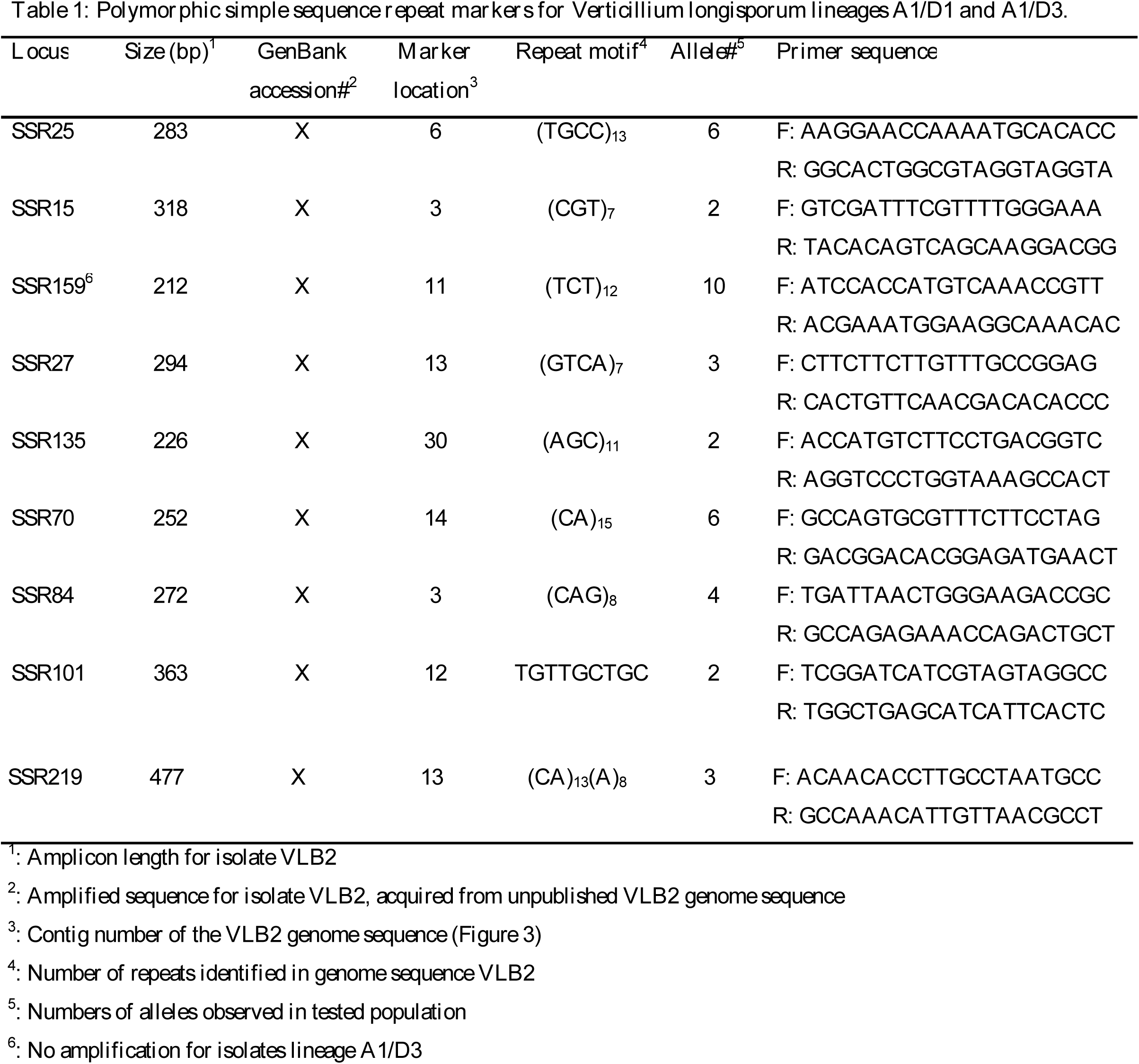
**Polymorphic simple sequence repeat markers for *Verticillium longisporum* lineages A1/D1 and A1/D3**.

The population structure was assessed based on the polymorphisms of these nine SSR loci that are dispersed over the genome (Table 1). The acquired multi-locus genotype (MLG) data were used to determine the most likely amount of genetic clusters in the *V. longisporum* population, allowing individual isolates to admix between different genetic clusters. The ad-hoc statistic Δ*K* was maximized for 3 genetic clusters in the population (K = 3), indicating that 3 is the most likely number of genetic clusters for the complete data set (Figure 1; Figure S1). Four previously characterized A1/D3 isolates formed one genetic cluster, whereas isolates known to belong to lineage A1/D1 showed two distinct clusters: one that includes samples from the USA and Japan, and another with samples from Germany and Sweden (Figure 1; Figure S1). The hitherto uncharacterised isolates all grouped together with one of these two A1/D1 clusters, indicating that they belong to the A1/D1 lineage (Figure 1; Figure S1). Indeed, successful amplification of the lineage A1/D1 specific primers D1f/AlfD1r (Inderbitzin *et al.* 2013) confirmed that these isolates belong to the A1/D1 lineage (Figure S2).

**Figure 1.**
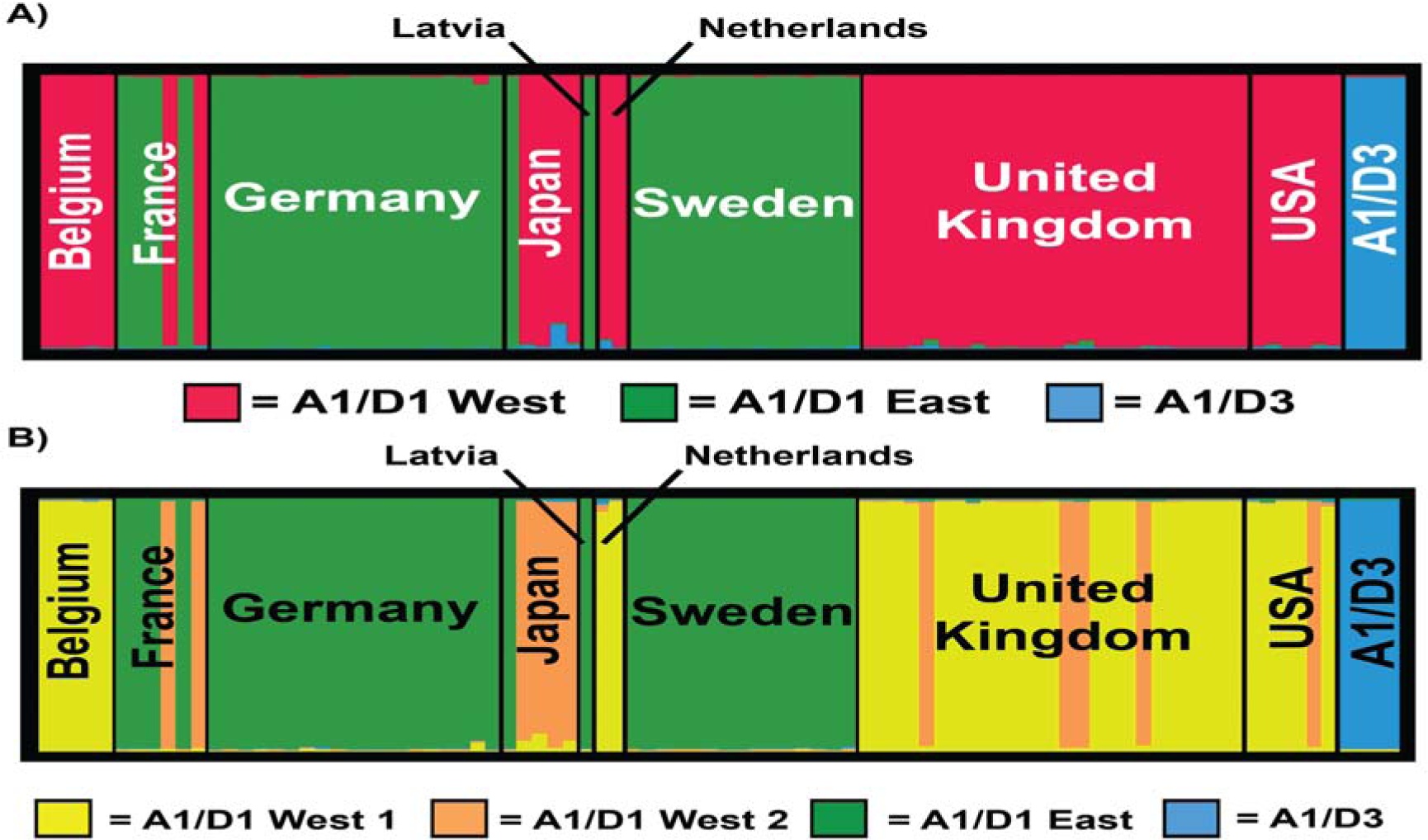
Clustering of individual *Verticillium longisporum* multi-locus genotypes (MLGs) using 15 polymorphic simple sequence repeat markers. Population clustering was executed on the whole data set for (A) three genetic clusters (K = 3) and (B) four genetic clusters (K = 4) with by the software Structure version 2.3 (Pritchard *et al.* 2000). The thick vertical bars separate the MLGs by country of origin. The bar width of every country is relative to the amount of samples: Belgium (*n* = 5), France (*n* = 6), Germany (*n* = 19), Japan (*n* = 5), Latvia (*n* = 1), the Netherlands (*n* = 2), Sweden (*n* = 15), UK (*n* = 25), USA (*n* = 6) and the cluster with isolates from the A1/D3 lineage (*n* = 4). The different colours represent separate genetic clusters. In A: red = lineage A1/D1 West, green = lineage A1/D1 East, and blue = lineage A1/D3. Lineage A1/D1 West is subdivided in section B, where yellow = A1/D1 West cluster 1, and orange = A1/D1 West cluster 2.

In addition, similar to all tested *V. longisporum* isolates, all uncharacterized isolates contained *MAT1-1* idiomorphs and failed to display *MAT1-2* in a mating type idiomorph PCR screen (Inderbitzin *et al.* 2011b) (Figure S3). Interestingly, the dichotomy within the A1/D1 lineage correlated with the country of origin of the isolates (*ρ* = -0.51, *p* = 0.00), as the Belgian, Dutch, UK and USA isolates formed one of the A1/D1 clusters (A1/D1 West; n = 44), whereas the German, Latvian and Swedish formed the other cluster (A1/D1 East; n = 40). These two clusters within lineage A1/D1 are further referred to as “lineage A1/D1 West” and “lineage A1/D1 East” according their relative geographic location in Europe. Furthermore, isolates from both genetic clusters are found in France and in Japan.

In order to confirm the dichotomy within lineage A1/D1 and to reveal more potential sub-structuring in the population, six additional published polymorphic SSR loci were used in the analysis (Table 2) (Barbara *et al.* 2005; Atallah *et al.* 2010). The SSR marker VDA817 from Barbara *et al.* (2005) also revealed polymorphisms, but was targeting the same SSR locus as VDA783 where CGTs are present in repetition. Thus, VDA817 was excluded from the population analysis. In total, 13 SSR loci were polymorphic for lineage A1/D1 as two of the nine previously mentioned SSR loci (SSR101 and SSR135) only differentiated between the A1/D1 and A1/D3 lineages (Table 1). The dichotomy within lineage A1/D1 was confirmed with these additional SSR loci, as the most likely number of genetic clusters was two for the A1/D1 population (Figure 1; Figure S3). Here, nine of the 13 polymorphic SSR loci for lineage A1/D1 subdivided the population significantly into the A1/D1 West and East dichotomy based on the analysis of molecular variance (AMOVA) (Φ_PT_ statistic, Table 2). Moreover, isolates of A1/D1 West and A1/D1 East lineages have no alleles in common for the loci SSR2511, SSR70, SSR219, VD8 and VDA823. All SSR loci combined, over 59% of the total genotypic variability within the A1/D1 population can be explained by the A1/D1 West and East dichotomy. Based on this, the dichotomy within A1/D1 is considered significant (*Φ*_*PT*_ = 0.590; *p* = 0.001).

**Table 2.** **The cluster determining capacities of the A1/D1 polymorphic simple sequence repeat loci**. Total number of alleles is given and the number of alleles that exclusively occur in A1/D1 West or A1D1 East. The *Φ_PT_* statistic and its significance were calculated for the SSR loci, regarding the 2 genetic clusters A1/D1 West and East.

| Locus | Marker<br>location<sup>1</sup> | Repeat motif<sup>2</sup> | Allele #<sup>3</sup> | A1/D1<br>West<sup>4</sup> | A1/D1<br>East<sup>5</sup> | $\Phi_{PT}$<sup>6</sup> | *p* |
| --- | --- | --- | --- | --- | --- | --- | --- |
| SSR2511 | 6 | (TGCC)<sub>13</sub> | 5 | 4 | 1 | 0.827 | 0.001 |
| SSR15 | 3 | (CGT)<sub>7</sub> | 2 | 0 | 0 | -0.005 | 0.468 |
| SSR159 | 11 | (TCT)<sub>12</sub> | 10 | 7 | 2 | 0.537 | 0.001 |
| SSR27 | 13 | (GTCA)<sub>7</sub> | 2 | 1 | 0 | -0.002 | 1.000 |
| SSR70 | 14 | (CA)<sub>15</sub> | 5 | 3 | 2 | 0.610 | 0.001 |
| SSR84 | 3 | (CAG)<sub>8</sub> | 3 | 2 | 0 | 0.006 | 0.492 |
| SSR219 | 13 | (CA)<sub>13</sub>(A)<sub>8</sub> | 2 | 1 | 1 | 1.000 | 0.001 |
| VD1 | 9 | (GCCT)<sub>8</sub><sup>7</sup> | 2 | 0 | 1 | 0.951 | 0.001 |
| VD8 | 11 | (TGA)<sub>13</sub> | 5 | 3 | 2 | 0.662 | 0.001 |
| VD12 | 6 | (CTTT)<sub>19</sub> | 9 | 1 | 1 | 0.267 | 0.002 |
| VDA783<sup>8</sup> | 6 | (CGT)<sub>22</sub> | 5 | 1 | 2 | 0.026 | 0.092 |
| VDA787 | 4 | (GAC(AGC)<sub>2</sub>)<sub>7</sub> | 2 | 1 | 0 | 0.735 | 0.001 |
| VDA823 | 5 | (CCG)<sub>8</sub> | 2 | 1 | 1 | 1.000 | 0.001 |

<sup>1</sup>: Contig number of the VLB2 genome sequence (Figure 3)
<sup>2</sup>: Number of repeats identified in genome sequence VLB2
<sup>3</sup>: Numbers of alleles observed in tested A1/D1 population
<sup>4</sup>: Number of alleles exclusive for A1/D1 West
<sup>5</sup>: Number of alleles exclusive for A1/D1 East
<sup>6</sup>: Computed with GenAlEx version 6.502 (Peakall & Smouse 2006, 2012)
<sup>7</sup>: Different repeat motif than reported for *Verticillium dahliae* in Atallah et al. (2010)
<sup>8</sup>: Targets same locus as VDA817 from Barbara *et al.* (2005)

Although our data divided the A1/D1 population into two genetic clusters, an additional population structure analysis was performed for lineage A1/D1 West and A1/D1 East separately to reveal putative sub-structuring. For the A1/D1 West population, the ad-hoc statistic Δ*K* was maximized for 2 genetic clusters in the population (K = 2), indicating that 2 is the most likely number of genetic clusters within the A1/D1 West cluster (Figure 1; Figure S4). Here, the Belgian and Dutch isolates formed one genetic cluster that segregated from the French/Japanese isolates. The UK and USA populations both resided in A1/D1 West clusters. In addition, AMOVA analysis also confirmed this sub-division within A1/D1 West as almost 34% of the genotypic variability within A1/D1 West can be explained by this dichotomy (*Φ*_*PT*_ = 0.338; *p* = 0.001). Further genetic clustering of the A1/D1 West population (K > 2) did not reveal any more subdivisions in A1/D1 West as isolates were then assigned to more than one genetic cluster. Furthermore, no further genetic clusters were present in lineage A1/D1 East as isolates were more or less equally subdivided between two clusters when K = 2. In conclusion, the *V. longisporum* lineage A1/D1 population contained 3 genetic clusters with no apparent intermixing between clusters, although intermixing between genetic clusters was enabled in the Structure analysis. The lack of intermixing between the genetic clusters within lineage A1/D1 indicates an exclusively clonal reproduction (Figure 1). The standardized index of association was calculated to investigate linkage disequilibrium between loci of *V. longisporum* lineage A1/D1. *I*^S^_A_ was significantly different from 0 (*I*^S^_A_ = 0.3081, *p* < 1.00 x 10^-5^) indicating that *V. longisporum* is not out-crossing.

The *V. longisporum* A1/D1 collection contained 40 different MLGs derived from 79 isolates with a complete genotype, of which 31 were unique (Table 3). The diversity between the UK isolates was higher than between the German and Swedish ones as Nei’s corrected gene diversity (*H*_*s*_) was 0.163, 0.110 and 0.085, respectively (Table 3). In agreement with this, the diversity of the whole A1/D1 West (*H*_*s*_ = 0.176) cluster was higher than A1/D1 East (*H*_*s*_ = 0.109). The difference in diversity between A1/D1 West and A1/D1 East is also clearly depicted in the genealogical network of the isolates; excluding locus VD12 to reduce the total amount of MLGs to 31 MLGs (Figure 2). The A1/D1 West and A1/D1 East populations were segregated from each other by a minimal of 10 mutations between MLG 3 and MLG 17. The A1/D1 East network was centred on the modal MLG 6 that represents more than half of the isolates (n=22) from four different countries (France, Germany, Sweden and Latvia). In contrast, A1/D1 West had a less centralized population network with MLGs 15, 23 and 26 being the most represented with 8, 9 and 7 individuals, respectively. MLG 15 contains exclusively UK isolates, whereas MLG23 and MLG26 have representatives from multiple countries (Belgium, Netherlands, UK and USA).

**Table 3:**
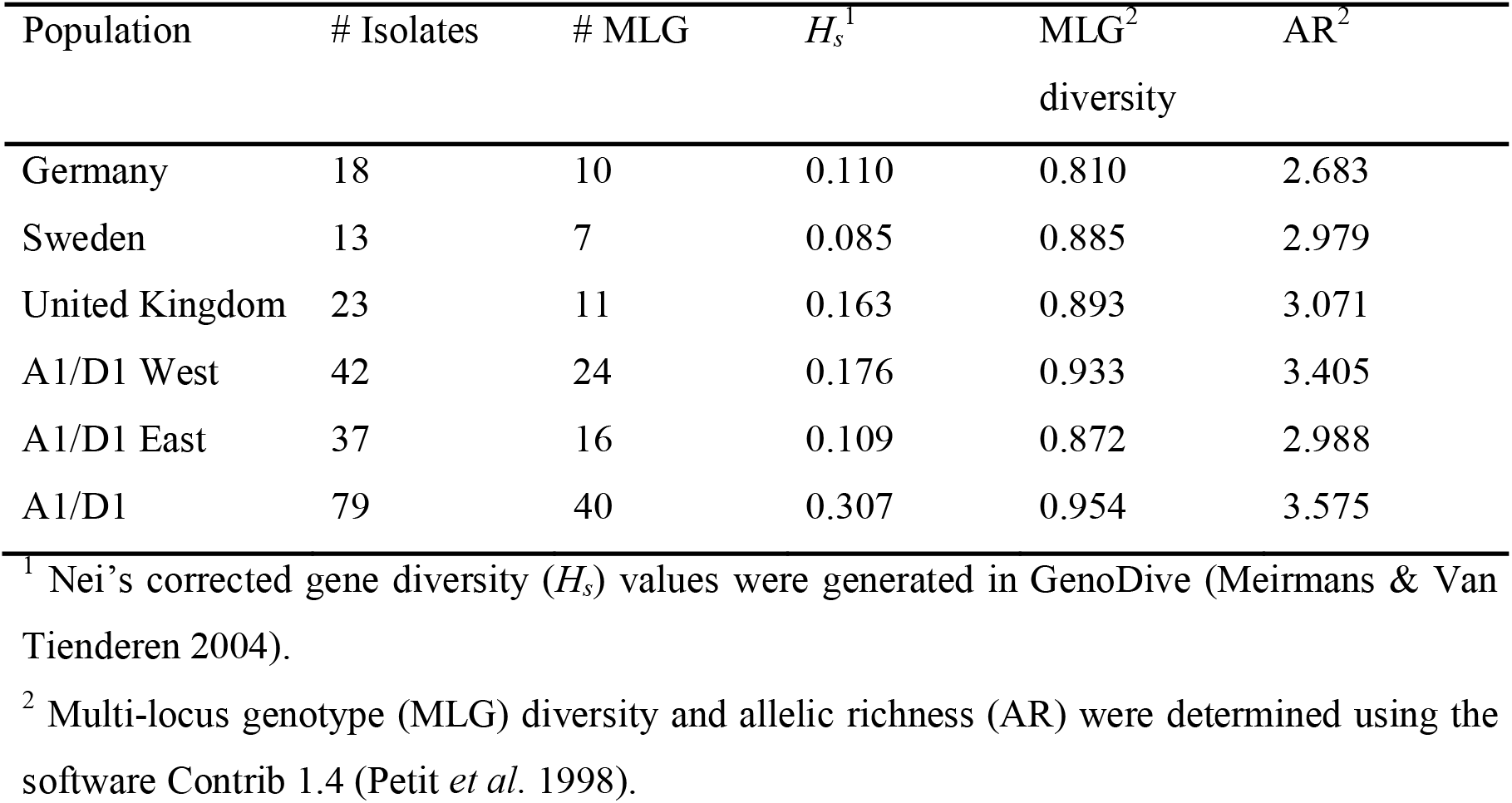
Diversity within the German, Swedish, UK, A1/D1 West, A1/D1 East and A1/D1 population.

**Figure 2.**
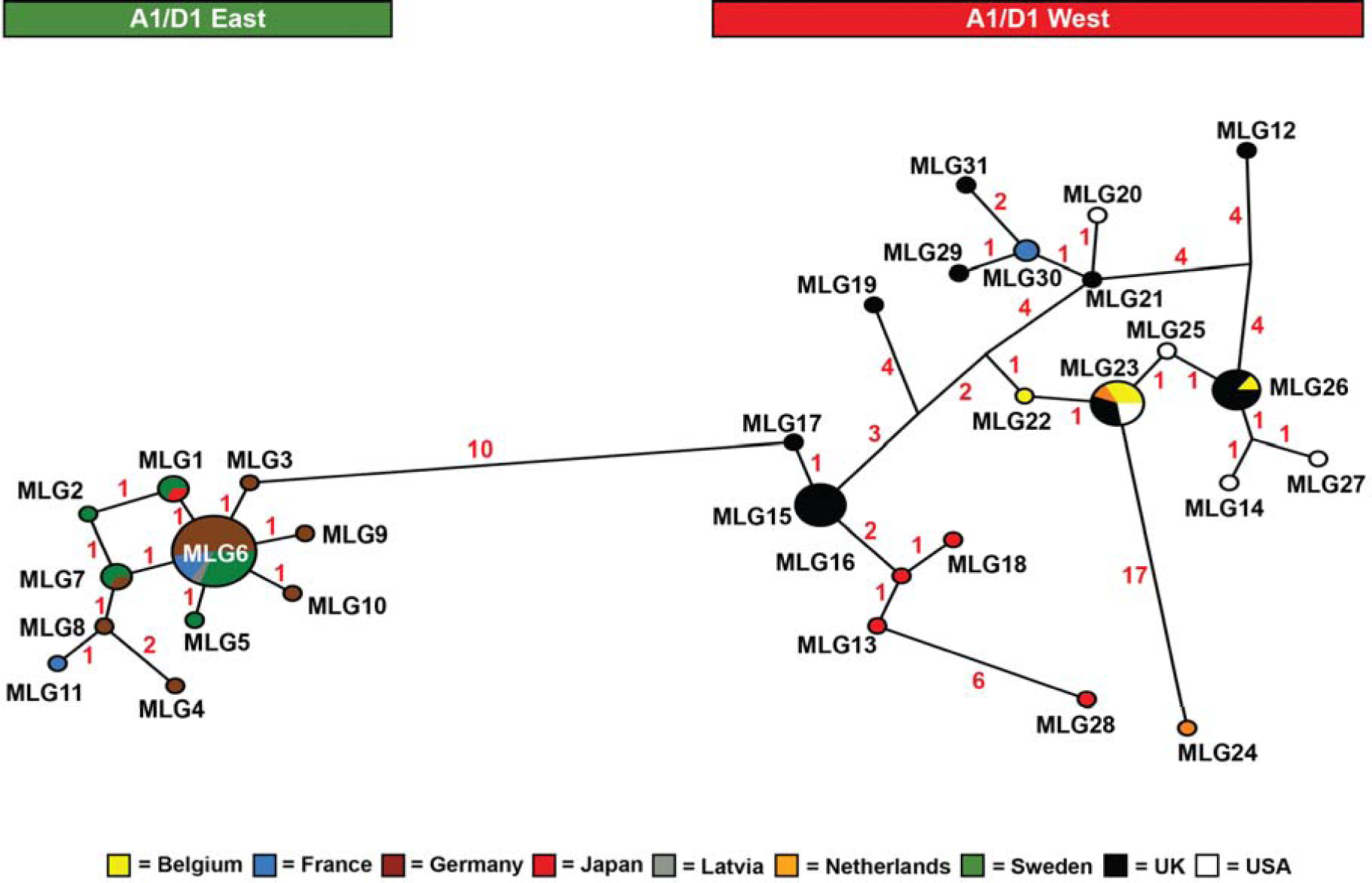
Geneological relationships of *Verticillium longisporum* A1/D1 population based on 12 polymorphic simple sequence repeat (SSR) loci. Geneological relationships were calculated based on 12 SSR loci (Table 2 except marker VD12) using the software Network 5.0.0.0 (Bandelt *et al.* 1999). The median-joining (MJ) network algorithm was chosen whereby SSR loci were weighted equally (10) and epsilon = 0 was used. Every circle represents a different multi-locus genotype (MLG) and the radius is relative to the MLG’s occurrence in the population. The lines connecting the MLGs depict the genealogical relationship between them, whereby the number of mutations between MLGs is written next to the lines. All MLGs on the left of the figure (under the green bar) clustered to the previous determined lineage A1/D1 East, whereas all the MLGs on the right (under the red bar) are members of the A1/D1 West cluster. MLGs with missing data were excluded.

The origin of the dichotomy within lineage A1/D1 (Figure 1) can either point to two independent hybridization events, giving rise to the two subpopulations within A1/D1, or to a single hybridization event followed by evolutionary diversification, leading to the emergence of two sub-populations. To gather additional evidence supporting either hypothesis, we selected one *V. longisporum* isolate of A1/D1 West (VL20) and one of A1/D1 East (VLB2) for single-molecule real-time (SMRT) sequencing using the PacBio RSII platform, which has been previously demonstrated to deliver high quality genome assemblies of *V. dahliae* and *Verticillium tricorpus* (Faino *et al.* 2015; Seidl *et al.* 2015). We generated 466,673 and 457,986 filtered subreads (~67x coverage) for *V. longisporum* strain VL20 and VLB2 that were assembled into 74 and 83 contigs, respectively (Table S2). Subsequent manual curation yielded a final assembly of 44 and 45 contigs with a total assembled genome length of 72.3 and 72.9 Mb for *V. longisporum* VL20 and VLB2, respectively (Table S2), which represents approximately twice the size of the recently assembled complete, telomere-to-telomere assemblies of *V. dahliae* strains JR2 (36.2 Mb) and VdLs17 (36.0 Mb) (Faino *et al.* 2015).

To place the sequenced *V. longisporum* strains into the context of other *Verticillium* species, we extracted the alleles of four protein-coding genes, namely actin (*ACT*), elongation factor 1-alpha (*EF*), glyceraldehyde-3-phosphate dehydrogenase (*GPD*) and tryptophan synthase (*TS*) (Inderbitzin *et al.* 2011b), from the two genome assemblies and performed maximum likelihood phylogenetic analyses (Figure S6), confirming that both *V. longisporum* isolates belong to the A1/D1 lineage.

In order to obtain further evidence for their evolutionary origin and to fully utilize the genome assemblies of the two *V. longisporum* isolates, we performed whole-genome comparisons between *V. longisporum* VL20 and VLB2 (Figure 3A). Whole-genome alignments between *V. longisporum* VL20 and VLB2 revealed large-scale synteny between both genomes, where a single genomic region in one of the genome aligns to two regions in the other genome. Moreover, as expected from an interspecific hybrid, the alignments of one of these regions generally displays >99% identity, while the identity of the second region is lower, ranging from 90 to 95% (Figure 3B). Notably, comparisons between *V. longisporum* strains VL20 and VLB2 only identified 140 kb and 450 kb, respectively, of genomic material that is absent in the other strain (Figure 3B). Furthermore, ~1,000 SNPs and ~3,800 indels between VL20 and VLB2 were revealed. Thus, the two *V. longisporum* strains are genetically highly similar and do not display marked differences between their individual LS regions; a likely scenario if the two *V. longisporum* strains emerged from the same hybridization event.

**Figure 3.**
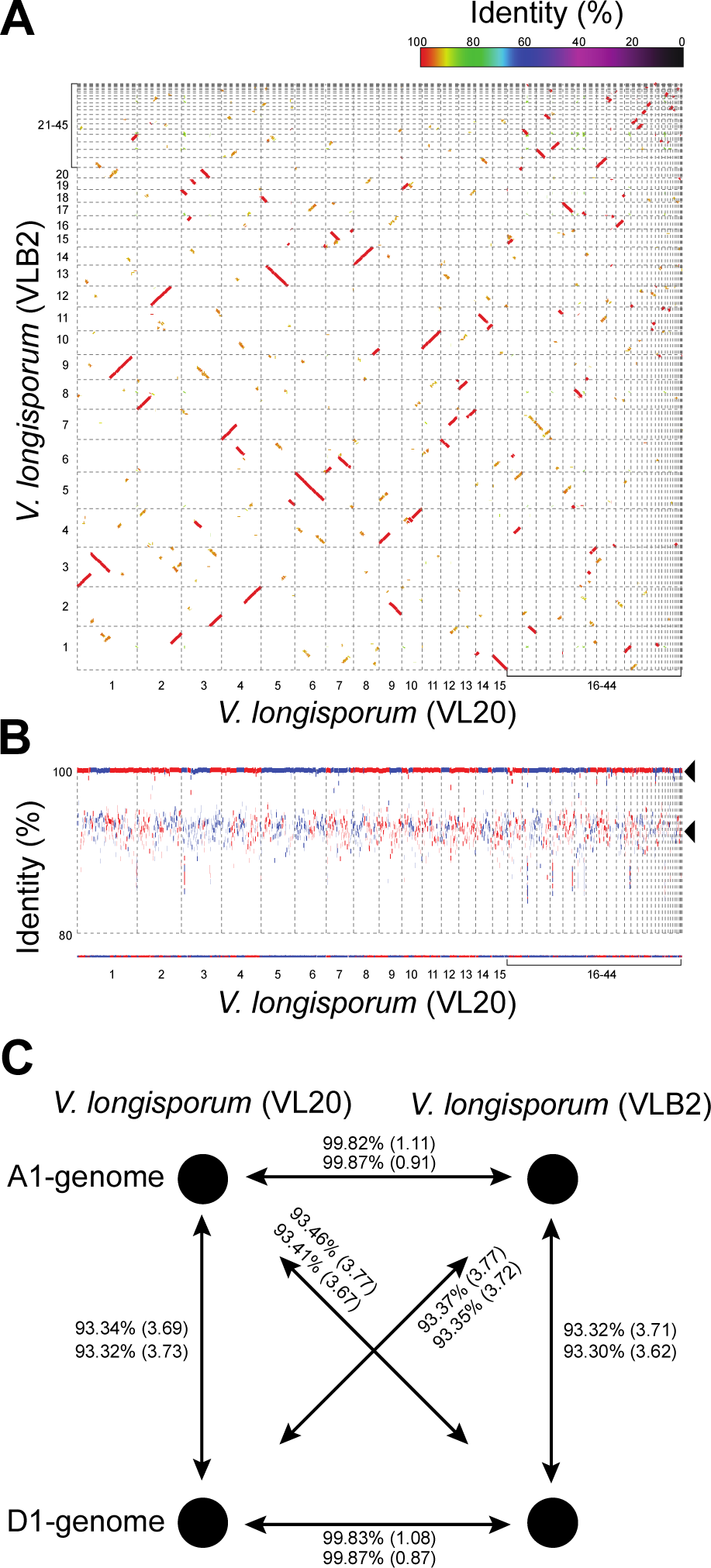
Whole-genome comparison between *Verticillium longisporum* VLB2 and VL20 reveals a common origin. A) Whole-genome comparision between *V. longisporum* VL20 and VLB2. Aligned genomic regions (>10 kb) are displayed and colour indicates sequence identity. B) Coverage plot displaying the whole-genome alignment (>10 kb) of *V. longisporum* strain VLB2 to that of strain VL20. Since *V. longisporum* is an allopolyploid, two genomic regions, one with ~99% and the other with 90-95% sequence identity (indicated by the two black arrow heads), align to the reference genome. Forward-forward alignments are shown in red and forward-reverse alignments (inversions) are shown in blue. C) Sequence identity between the A1 and D1 sub-genomes of *V. longisporum* strains VLB2 and VL20.

To obtain further evidence the evolutionary origin of the two *V. longisporum* strain, we extracted the individual A1 and D1 sub-genomes based on their sequence identity to *V.dahliae* stain JR2, as parent D1 is presumed to belong to *V. dahliae* (Figure S6) (Inderbitzin *et al.* 2011b). Moreover, genome comparisons between *V. dahliae* and *V. longisporum* indicated that 95% sequence identity allows discriminating A from D sub-genome. The extracted A1 and D1 sub-genomes of *V. longisporum* strain VL20 comprised 32.8 Mb and 34.2 Mb, respectively, while 5.2 Mb could not be assigned. Similarly, the A1 and D1 sub-genomes of *V. longisporum* strain VLB2 comprised 32.5 Mb and 34.7 Mb, respectively, while 5.7 Mb could not be assigned. As expected, within and between the *V. longisporum* strains the average identity between A1 and D1 sub-genomes was ~93% (Figure 3C). Notably, however, the average identity between the A1 sub-genomes as well as between the D1 sub-genomes of both *V. longisporum* strains was >99% (Figure 3C), suggesting that the A1 and D1 parental genomes that give rise to *V. longisporum* strain VL20 and VLB2 were identical and that, consequently, A1/D1 West and A1/D1 East are derived from a single hybridization event (Figure 4).

**Figure 4.**
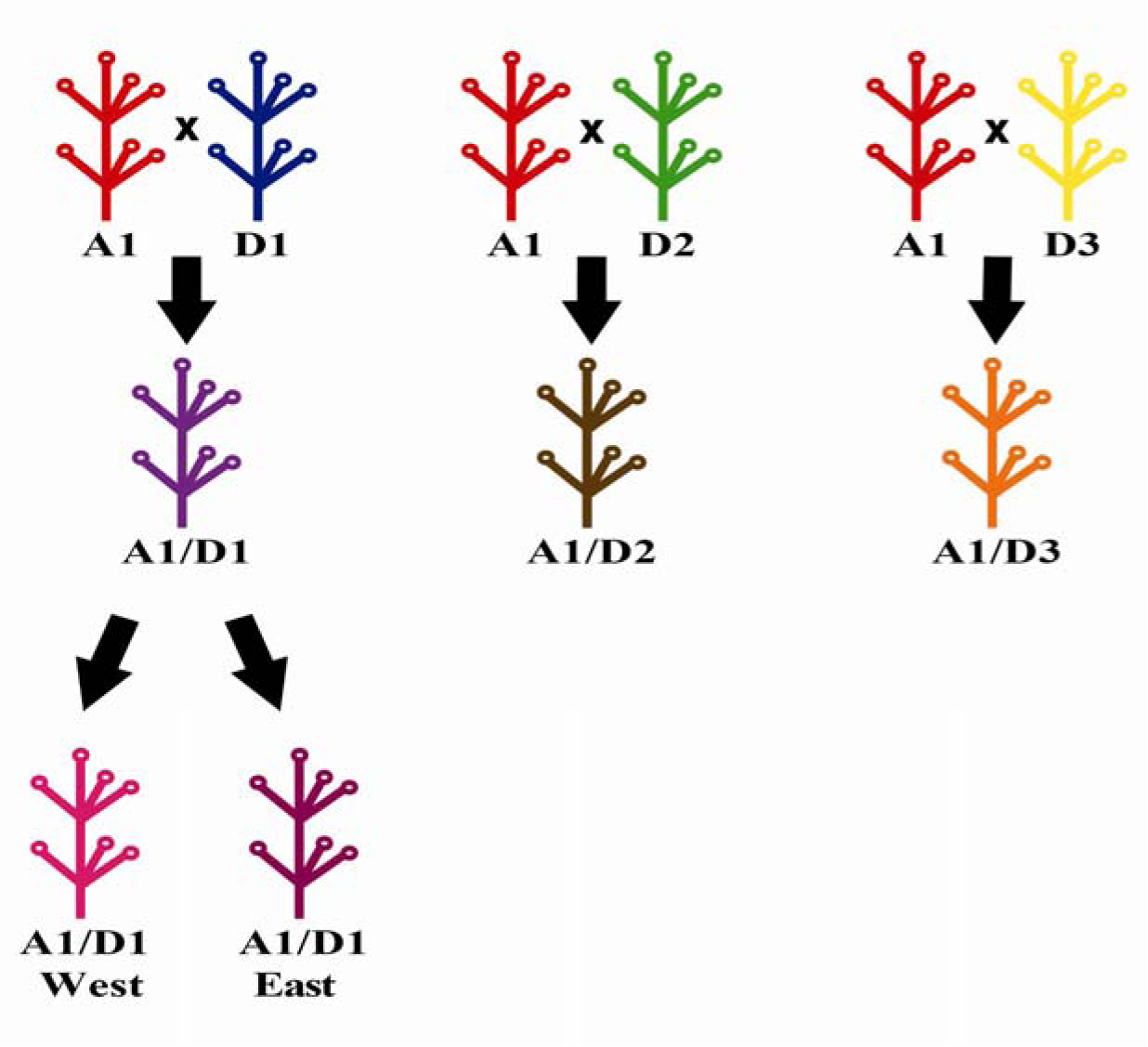
The evolutionary history of *Verticillium longisporum*. *V. longisporum* consists of three seperate hybridization events between two different *Verticillium* spp. Hybrid lineages are named after their parental species: A1/D1, A1/D2 and A1/D3. Two A1/D1 populations have segregated from each other fur still unclear reasons. Phylogenetic relationships between parents of *V. longisporum* and other *Verticillium* spp. are depicted in figure S6. (Adjusted from Depotter *et al.*, 2016a).

## Discussion

Population genetic structures reveal information about the evolutionary history of organisms. Genome hybridisation can be a major driver for organismal adaptation and has allowed *V.dahliae* that infect brassicaceous species relatively infrequently, to become pathogenic on these hosts as *V. longisporum* (Depotter *et al.* 2016a). Based on phylogenetic analyses, *V. longisporum* has been subdivided into three lineages: A1/D1, A1/D2 and A1/D3, each representing a separate hybridization event between two *Verticillium* species (Inderbitzin *et al.* 2011b). The Verticillium stem striping pathogen has been emerging as a disease on oilseed rape and is emerging in hitherto unaffected production regions (Gladders *et al.* 2011; CFIA 2015). This disease is predominantly caused by the *V. longisporum* lineage A1/D1, which is the most virulent lineage on this crop (Novakazi *et al.* 2015). In this study, the diversity within the A1/D1 lineage was assessed and population structures within a collection from a diverse geographic origin were elucidated. Isolates from nine different countries (Table S1) were genotyped with newly and previously characterized polymorphic SSR loci (Tables 1&2). Model-based clustering revealed a hitherto undiscovered dichotomous structuring within *V. longisporum* lineage A1/D1 (Figure 1; Figure S1). Interestingly, the two A1/D1 sub-populations were geographically correlated and were accordingly labelled “A1/D1 West” and “A1/D1 East” based on their relative European location. Geographic population structuring typically indicates adaptation to local climate or to local hosts. Alternatively, the two genetic clusters may also represent two separate hybridization events. In order to test the latter hypothesis, whole-genome comparisons between two representative genomes of the A1/D1 West and East population were performed. All our evidence points towards a single origin of the two A1/D1 populations (Figure 4). First of all, the genome sizes of strains VL20 and VLB2 are highly similar, with 72.3 Mb and 72.9 Mb, respectively (Table S2). Then, the two genomes carried only a small propostion of lineage specific sequence: 140 kb and 450 kb for VL20 and VLB2, respectively. Recently, genome-comparisons between multiple *V. dahliae* strains have revealed the presence of extensive (2.5-4.5 Mb) lineage-specific (LS) regions that are shared by only a subset of the *V. dahliae* isolates and that enriched for *in planta* induced genes that contribute to fungal virulence (de Jonge *et al.* 2013; Faino *et al.* 2016). For example, direct comparisons between the completely assembled genomes of *V. dahliae* strains JR2 and VdLS17 (Faino *et al.* 2015), two strains that have been shown to be extremely closely related with 99.98% sequence identity (de Jonge *et al.* 2013), identified four large regions comprising ~2 Mb that display frequent presence/absence polymorphisms (Faino *et al.* 2016), of which in total ~550 kb and ~620 kb in *V. dahliae* strain JR2 and VdLs17 do not align to the other strain, respectively. Additionally, genome-wide comparison between *V.dahliae* strains JR2 and VdLs17 revealed ~4,700 SNPs and ~10,000 indels, while the *V. longisporum* VL20 and VLB2 strains contain genomes that were relatively low in divergence as only ~1,000 SNPs and ~3,800 indels were found. Finally, the genetic distance between the A sub-genomes as well as between the D-sub-genomes of both *V. longisporum* strains was >99% (Figure 3C), suggesting that the A and D parental genomes were nearly identical. Thus, we conclude that the observed geographic population structuring is a signature of divergent evolution driven by environmental adaptation.

Presently, environmental conditions responsible for the dichotomous A1/D1 population structure remain elusive. However, strikingly, UK A1/D1 isolates are more diverse than the German, Swedish and even the whole A1/D1 East population (Table 3). High diversity is unexpected for a disease that is considered to have been introduced recently (Gladders *et al.* 2011), especially regarding the isolating character of the British island (Nei *et al.* 1975). In contrast, Verticillium stem striping is an established disease in Germany and Sweden, with disease reports appearing since the 1960s (Stark 1961; Svensson & Lerennius 1987). Thus, the genetically diverse UK population disagrees with a recent introduction (Gladders *et al.* 2011), as introductions are considered a population bottleneck with a drastic loss of genetic variation. Potentially, genetic diversity may also be the consequence of multiple introductions. For example, three genetic lineages of the chestnut blight fungus *Cryphonectria parasitica* were found to be separately introduced in France (Dutech *et al.* 2010). However, the UK population consists of 11 MLGs (Table 3) that are genealogically widely dispersed among the A1/D1 West clade (Figure 2). Taking into account the novelty of the disease in the UK and, thus, the short timeframe of pathogen evolution, the high diversity is conceivably not caused by multiple introductions. Rather, the A1/D1 West population must have been present in the UK for a longer time without causing Verticillium stem striping on oilseed rape. In support of this, *V. longisporum* reported to be present already since 1957 in the UK, albeit not on oilseed rape but on Brussels sprouts (Isaac 1957; Karapapa *et al.* 1997).

The recent emergence of Verticillium stem striping combined with the high diversity of the UK isolates is puzzling. The A1/D1 West sub-population, partly consisting of UK isolates, is more diverse than its A1/D1 East counterpart (Table 3, Figure 2). Their shared origin along with this difference in diversity indicates that A1/D1 East is a founder population of the originating A1/D1 West population. The more recent origin of A1/D1 East is also depicted in the genealogical network (Figure 2). A1/D1 East has a clear modal MLG (MLG6) with all other A1/D1 East MLGs centred on it. In contrast, the A1/D1 West population lacks a dominating genotype. A1/D1 East may have evolved from A1/D1 West in order to adapt to different climate conditions, hence the geographic correlation of the two A1/D1 sub-populations. However, arguably, climate difference between the European countries from this study (Belgium, the Netherlands, France, Germany, Latvia, Sweden and UK) are minimal as they are generally classified under the same climate zone (Kottek *et al.* 2006). Moreover, the USA and Japanese isolates followed the same A1/D1 West and East dichotomy, although climatological differences with Europe are expected to be bigger. Population segregation in organisms with symbiotic relationships can also be host driven. Genetic co-structuring between pathogen and host can be found due to their co-evolutionary interaction (Croll & Laine 2016; Feurtey *et al.* 2016), as pathogens engage in arm races with hosts for continued symbiosis (Cook *et al.* 2015). Host-pathogen evolution may explain the discrepancy in population diversity and the Verticillium stem striping emergences of the two A1/D1 sub-populations. Conceivably, A1/D1 East was initially the sole causal agent of Verticillium stem striping as the disease flourished in Sweden and Germany since the 1960s, countries with an exclusive A1/D1 East presence (Figure 1) (Kroeker 1970). This initial evolutionary advantage of A1/D1 East may have facilitated its establishment in North-central Europe. Recently, A1/D1 West emerged as a pathogen on oilseed rape as diseased oilseed rape stems from the UK belonged to this genetic cluster (Figure 1) (Gladders *et al.* 2011). Oilseed rape must have become susceptible to A1/D1 West for reasons that presently remain unclear. Pathogen adaptation may have led to the emergence of the disease. However, the large genetic diversity in the recently emerged UK Verticillium stem striping population indicates alternative hypotheses, as resistance breaking generally emerges from a single genotype (Figure 2, Table 5). In addition, the small time-lag between the first report and omnipresence of Verticillium stem striping in the UK of this largely immobile pathogen conforms this idea (Gladders *et al.* 2013). Alterations in environmental conditions (e.g. global warming, Siebold and Tiedemann, 2012) and oilseed rape cultivars are therefore more likely hypotheses for the sudden rise of the genomic diverse UK Verticillium stem striping population. Oilseed rape is a relatively new crop in the UK that was virtually unknown in the 1970 but is currently one of the main arable crops with approximate production area of 700,000 hectares (Wood *et al.* 2013). Intensive breeding efforts have been made to create a broad diversity of oilseed rape oil, such as the modern oo (canola) type (Wittkop *et al.* 2009). The novelty of Verticillium stem striping makes that cultivar resistance for this disease was not selected for, hence the possibility that more susceptible oilseed rape varieties have been commercialized over time.

*V. longisporum* is an interspecific hybrid between two separate *Verticillium* species (Inderbitzin *et al.* 2011b; Depotter *et al.* 2016a). Interspecific hybrids are regularly found to be impaired in their sexual reproduction (Greig 2009; Bertier *et al.* 2013), although this should be of little significance for *V. longisporum* as a sexual stage has not been described for any of the *Verticillium* species (Short *et al.* 2014). However, mating types, meiosis-specific genes and genomic recombination between clonal lineages have been observed for *V. dahliae* (Milgroom *et al.* 2014; Short *et al.* 2014). This suggests that *V. dahliae* may have cryptic or ancestral sexual reproduction. In contrast to population structure studies with *V. dahliae* (Atallah *et al.* 2010, 2012), no apparent intermixing between genetic clusters was observed for *V. longisporum*. Moreover, the was also significantly different from 0 for the *V. longisporum* lineage A1/D1 (*I*^S^_A_ = 0.3081, *p* < 1.00 x 10^-5^), which implies that no linkage equilibrium is present. These data indicate that *V. longisporum* reproduces exclusively in a clonal fashion and has never experienced sexual reproduction.

Increasing evolutionary ecology knowledge of a pathogen can play a pivotal role in disease management in order to protect ecosystems (Williams 2010). Population genetic studies are therefore often used to identify origins of pathogen introductions and adaptation pathways (Dutech *et al.* 2012; Gross *et al.* 2014). For example, the Italian introduction site of the North American forest pathogen *Heterobasidion irregulare* coincided with an American military base in World War II (Garbelotto *et al.* 2013). However, not all emerging diseases are preceded by a clearly identified recent introduction. Our study exemplifies that new diseases can emerge from a latent, previously established, microbial population. Thus, factors like weather and farming techniques may also lead to the emergence of novel disease, especially for fungal pathogens (Anderson *et al.* 2004). Besides, the two *V. longisporum* populations are generally still geographically bound to specific geographic locations in Europe. The driving forces causing the dichotomous population structure remain elusive, but conceivable differences in pathogenic traits may have contributed to the West/East segregation. A cautious approach is therefore appropriate to prevent further spread of the two populations, as de expansion of the population into new geographic regions may have unpredicted outcomes.

## Acknowledgements

The authors would like to thank the Marie Curie Actions programme of the European Commission that financially supports the research investigating the threat of *V. longisporum* to UK oilseed rape production. Work in the laboratories of B.P.H.J.T. and M.F.S is supported by the Research Council Earth and Life Sciences (ALW) of the Netherlands Organization of Scientific Research (NWO). We greatly appreciate all donors of *V. longisporum* isolates mentioned in Table S1.

## Data Accessibility

All newly characterized SSR loci have been deposited in X by the accession numbers (X-X).

## Author Contributions

The study was conceived by T.A.W. and B.P.H.J.T. and designed by J.R.L.D and M.F.S under supervision of B.P.H.J.T and T.A.W. Experiments were performed by J.R.L.D., M.F.S. and G.C.M.B. Data analysis and interpretation by J.R.L.D., M.F.S., B.P.H.J.T and T.A.W. The manuscript was written by J.R.L.D, M.F.S. and B.P.H.J.T. All authors read and approved the final manuscript.

## Tables and Figures

**Table S1.**
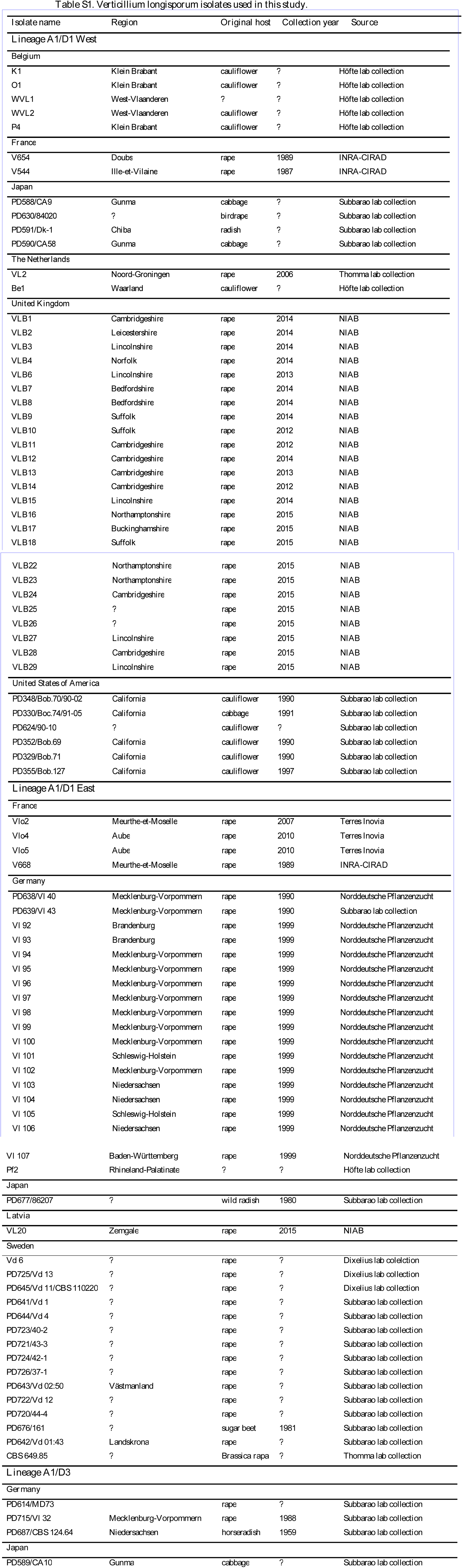
*Verticillium longisporum* isolates used in this study.

**Table S2:** *Verticillium longisporum* strain VL20 and VLB2 genome assemblies

| **Matric:** |  | **VL20** | **VLB2** |
| --- | --- | --- | --- |
| SMRT cells |  | 6 | 6 |
| Filtered subreads |  | 466,673 | 457,986 |
| Coverage (n-fold) |  | 67.7x | 66.4x |
| HGAP3 assembly | Size (bp) | 72,495,102 | 73,489,504 |
|  | Contigs | 74 | 83 |
|  | Longest contig | 5,837,573 | 5,411,788 |
|  | N<sub>50</sub> (bp) | 2,718,091 | 2,899,830 |
|  | No of Ns/100 kb | 0 | 0 |
| FinisherSC assembly | Size (bp) | 72,433,856 | 73,098,666 |
|  | Contigs | 52 | 56 |
|  | Longest contig | 7,148,864 | 5,411,733 |
|  | N<sub>50</sub> (bp) | 2,718,187 | 3,135,754 |
|  | No of Ns/100 kb | 0 | 0 |
| Final assembly | Size (bp) | 72,271,822 | 72,894,593 |
|  | Contigs | 44 | 45 |
|  | Longest contig | 7,148,928 | 5,411,750 |
|  | N<sub>50</sub> (bp) | 2,718,243 | 3,135,757 |
|  | No of Ns/100 kb | 0 | 0 |

**Figure S1.**
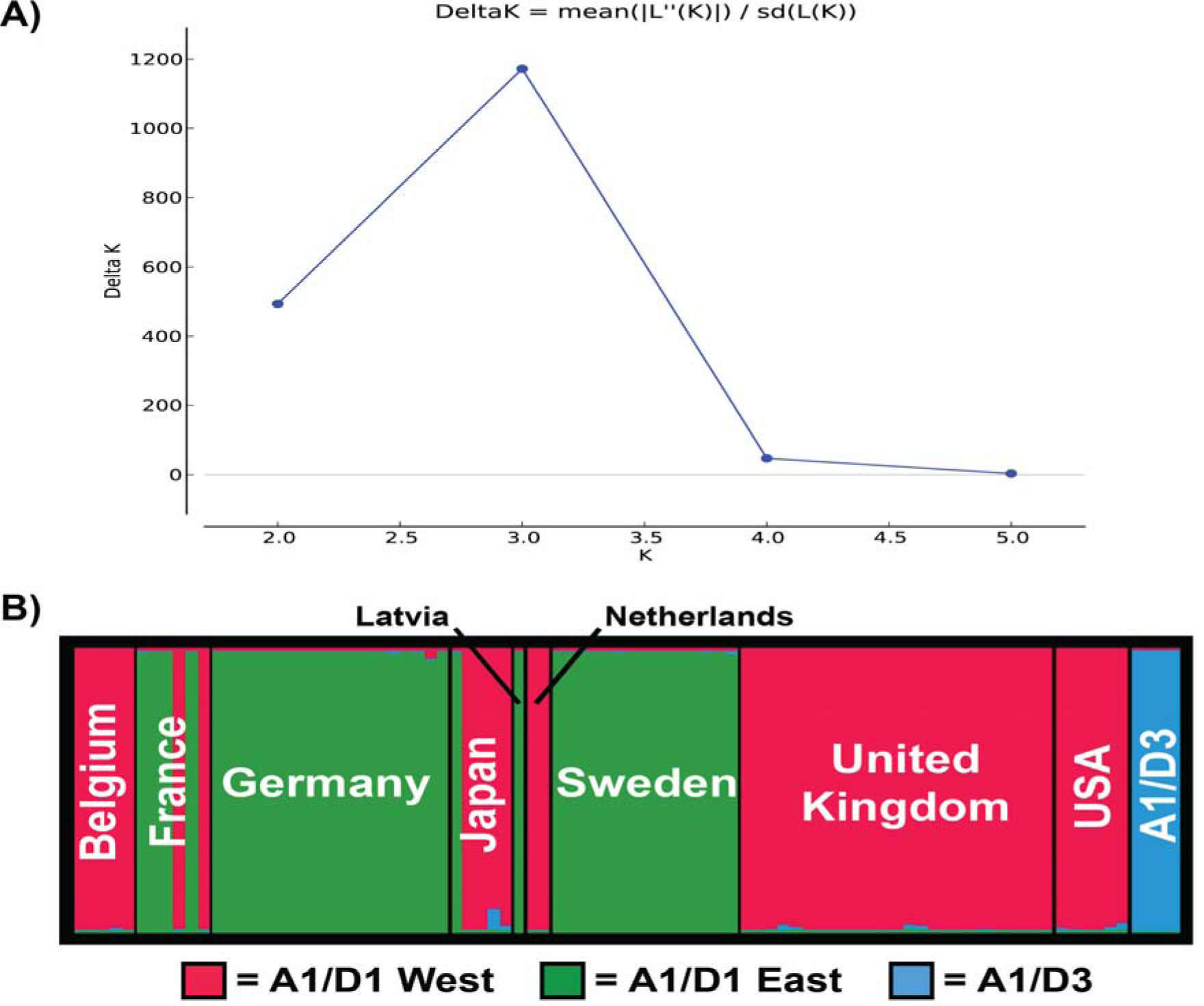
Genetic clusters within the *Verticillium longisporum* population using the 9 newly designed polymorphic simple sequence repeat (SSR) markers. Isolate genotyping was based on 9 SSR loci (Table 1). (A) Output of the ad-hoc statistic Δ*K* calculated for the different genetic clusters within *V. longisporum* population with a maximum for K = 3 (Earl & vonHoldt 2012). (B) Genetic clustering of individual *V. longisporum* multi-locus genotypes (MLGs) devided the whole data set into three groups using Structure version 2.3 (Pritchard *et al.* 2000). The thick vertical bars separate the MLGs by country of origin. The bar width of every country is relative to the amount of samples: Belgium (*n* = 5), France (*n* = 6), Germany (*n* = 19), Japan (*n* = 5), Latvia (*n* = 1), the Netherlands (*n* = 2), Sweden (*n* = 15), UK (*n* = 25), the USA (*n* = 6) and the cluster with isolates from the A1/D3 lineage (*n* = 4). The different colours represent separate genetic clusters: red = lineage A1/D1 West, green = lineage A1/D1 East, and blue = lineage A1/D3.

**Figure S2.**
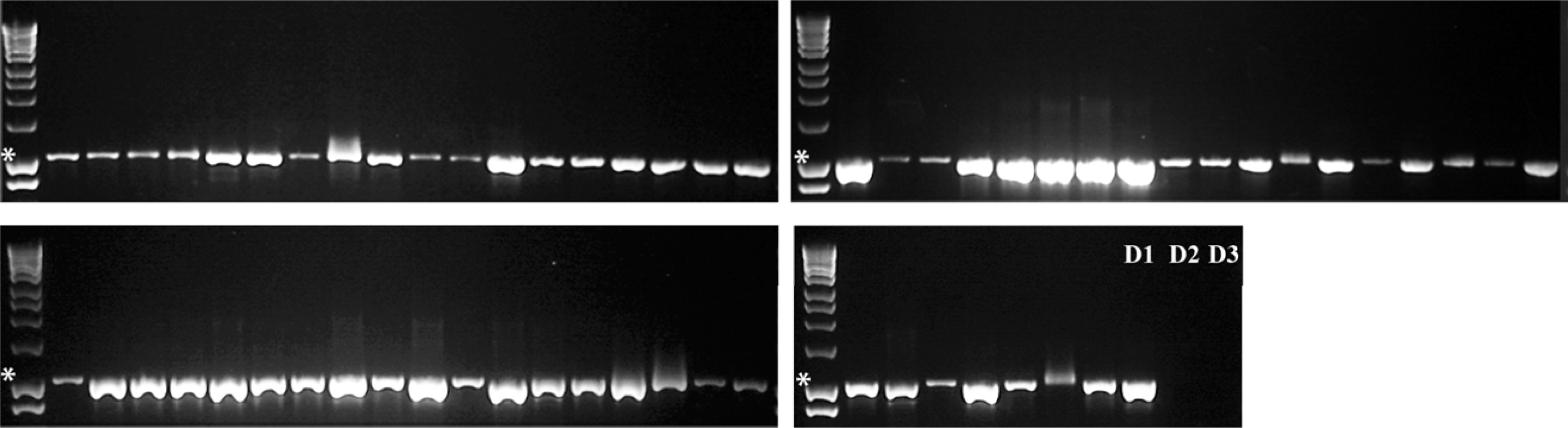
Lineage characterization of *Verticillium longisporum* isolates. Agarose gels show the selective amplification of markers obtained with the lineage A1/D1 specific primers D1f/AlfD1r (Inderbitzin *et al.* 2013) for hitherto uncharacterized *V. longisporum* isolates. Gels are deliminated by ladders where the 1000 bp size marker is indicated by by ‘*’. Isolates with an 1020 bp amplicon belong to the A1/D1 lineage. The three lanes indicated by ‘D1’, ‘D2’ and ‘D3’ are controls, in which previously characterized isolates were used from lineage A1/D1, A1/D2 and A1/D3 respectively. Isolates used from top left to bottom right: K1, O1, WVL1, WVL2, P4, Vlo2, Vlo4, Vlo5, V654, V668, V544, Pf2, Vl 40, Vl 98, Vl 99, Vl 100, Vl 101, Vl 102, Vl 103, Vl 104, Vl 105, Vl 106, Vl 107, Vl 43, PD677, VL20, Bel, VLB3, VLB4, VLB6, VLB7, VLB8, VLB9, VLB10, VLB11, VLB12, VLB13, VLB14, VLB15, VLB16. VLB17, VLB18, VLB22, VLB23, VLB24, VLB25, VLB26, VLB27, VLB28, VLB29, Vd 11, PD356 and PD715.

**Figure S3.**
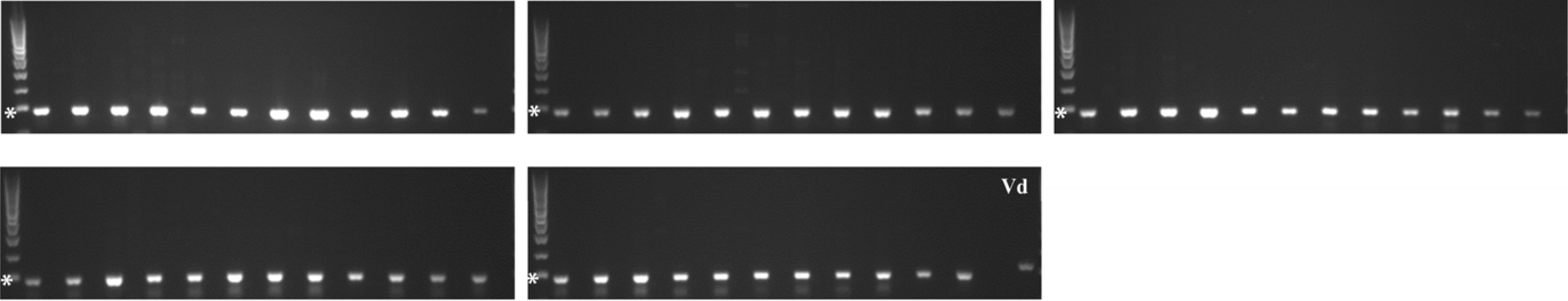
Mating type characterization of *Verticillium longisporum* isolates. Agarose gels shows alternatingly the presence/absence of the mating type idiomorphs *MAT1-1* and *MAT1-2* according to Inderbitzin *et al.* (2011b) for hitherto uncharacterized *V. longisporum* isolates. Gels are the 300 bp size marker is indicated by ‘*’. The *MAT1-1* and *MAT1-2* specific amplicons have a length of 291 and 330bp, respectively. The two lanes function as a positive control for *MAT1-2* as *Verticillium dahliae* (Vd) DNA of isolate JR2 was used as template. Isolates used from top left to bottom right: K1, O1, WVL1, WVL2, P4, Vlo2, Vlo4, Vlo5, V654, V668, V544, Pf2, Vl 92, Vl 93, Vl 94, 99, Vl 100, Vl 101, Vl 102, Vl 103, Vl 104, Vl 105, Vl 106, Vl 107, Vl 43, PD677, VL20, Be1, VL2, CBS 649.85, vd 6, VLB1, VLB2, VLB3, VLB4, VLB6, VLB7, VLB8, VLB9, VLB10, VLB11, VLB12, VLB13, VLB14, VLB15, VLB16, VLB17, VLB18, VLB22, VLB23, VLB24, VLB25, VLB26, VLB27, VLB28, VLB29 and JR2.

**Figure S4.**
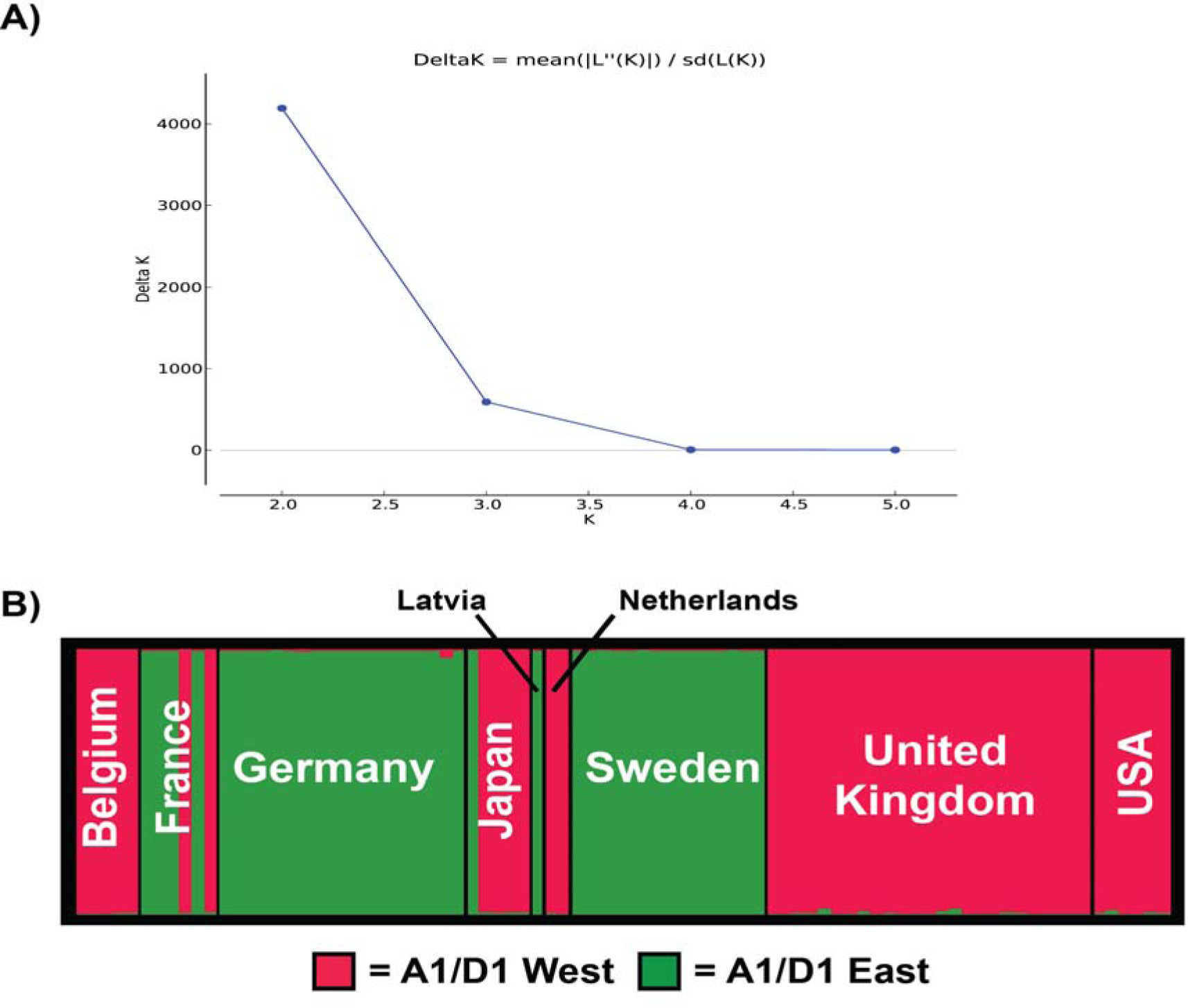
Genetic clusters within the *Verticillium longisporum* A1/D1 population using 13 polymorphic simple sequence repeat (SSR) markers. Isolate genotyping was based on 13 SSR loci (Table 2). (A) Output of the ad-hoc statistic Δ*K* calculated for the different genetic clusters within *V. longisporum* population with a maximum for K = 2 (Earl & vonHoldt 2012). (B) Genetic clustering of individual *V. longisporum* multi-locus genotypes (MLGs) into 2 groups determined by Structure version 2.3 (Pritchard *et al.* 2000). The thick vertical bars separate the MLGs by country of origin. The bar width of every country is relative to the amount of samples: Belgium (*n* = 5), France (*n* = 2), Japan (*n* = 5), the Netherlands (*n* = 2), UK (*n* = 25) and USA (*n* = 6). The different colours represent separate genetic clusters: red = A1/D1 West cluster and green = A1/D1 East cluster.

**Figure S5.**
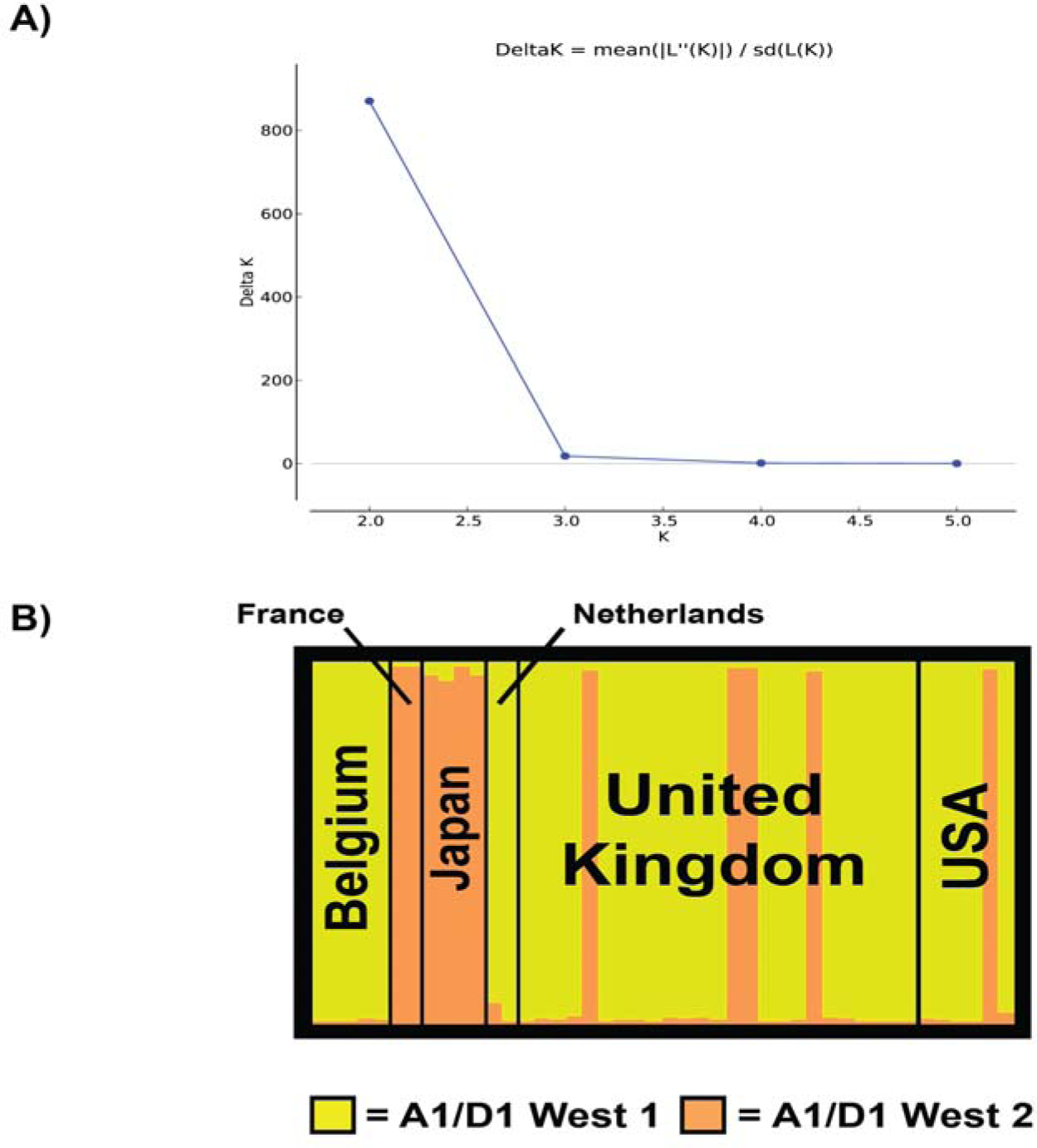
Genetic clusters within the *Verticillium longisporum* A1/D1 West population using 11 polymorphic simple sequence repeat (SSR) markers. Isolate genotyping was based on 11 SSR loci (Table 2 except markers SSR219 and VDA823). (A) Output of the ad-hoc statistic Δ*K* calculated for the different genetic clusters within *V. longisporum* population with a maximum for K = 2 (Earl & vonHoldt 2012). (B) Genetic clustering of individual *V. longisporum* multi-locus genotypes (MLGs) into 2 groups determined by Structure version2.3 (Pritchard *et al.* 2000). The thick vertical bars separate the MLGs by country of origin. The bar with of every country is relative to the amount of samples: Belgium (*n* = 5), France (*n =* 2), Japan (*n* = 4), the Netherlands (*n* = 2), UK (*n* = 25) and USA (*n* = 6). The different colours represent separate genetic clusters: yellow = A1/D1 West cluster 1 and orange green = A1/D1 West cluster 2.

**Figure S6.**
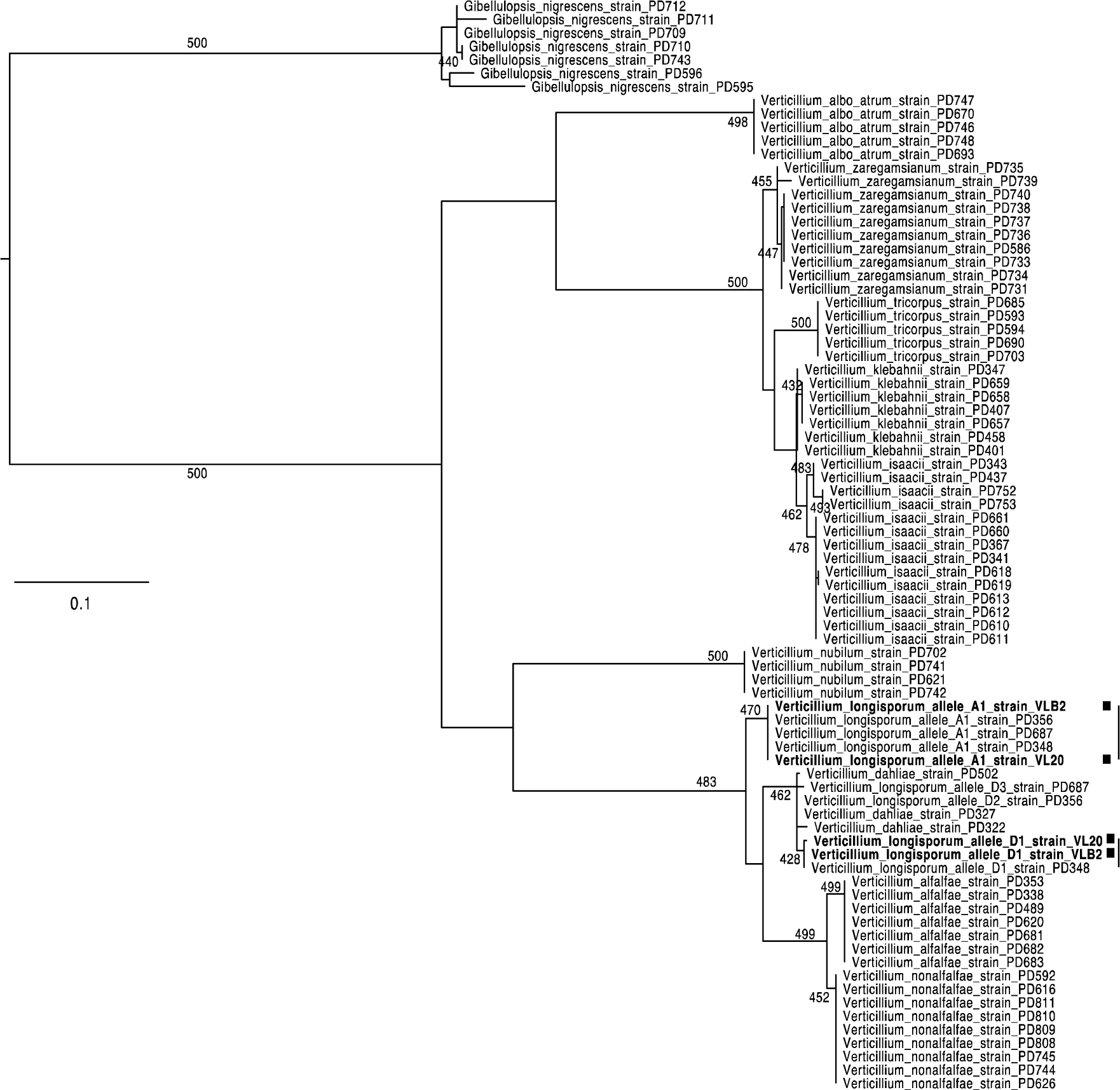
Phylogenetic relationships between *Verticillium longisporum* A1/D1 West and East parents with the other *Verticillium* species. Phylogenetic relationships are based on the sequence of four previously used protein-coding genes: actin (*ACT*), elongation factor 1-alpha (*EF*), glyceraldehyde-3-phosphate dehydrogenase (*GDP*) and tryptophan synthase (*TS*) (Inderbitzin *et al.* 2011a). The phylogenetic tree was reconstructed using PhyML using the GTR nucleotide substitution model and four discrete gamma categories (Guindon & Gascuel 2003). The robustness of the phylogeny was assessed using 500 bootstrap replicates. VLB2 and VL20 were used as a representative for the lineage A1/D1 West and East, respectively (Table S1). These were included together with 74 previously characterized *Verticillium* and *Verticillium*-related isolates in the phylogenetic analysis.

